# Dual Near Infrared Two-Photon Microscopy for Deep-Tissue Dopamine Nanosensor Imaging

**DOI:** 10.1101/145912

**Authors:** Jackson T. Del Bonis-O’Donnell, Ralph H. Page, Abraham G. Beyene, Eric G. Tindall, Ian McFarlane, Markita P. Landry

## Abstract

A key limitation for achieving deep imaging in biological structures lies in photon absorption and scattering leading to attenuation of fluorescence. In particular, neurotransmitter imaging is challenging in the biologically-relevant context of the intact brain, for which photons must traverse the cranium, skin and bone. Thus, fluorescence imaging is limited to the surface cortical layers of the brain, only achievable with craniotomy. Herein, we describe optimal excitation and emission wavelengths for through-cranium imaging, and demonstrate that near-infrared emissive nanosensors can be photoexcited using a two-photon 1560 nm excitation source. Dopamine-sensitive nanosensors can undergo two-photon excitation, and provide chirality-dependent responses selective for dopamine with fluorescent turn-on responses varying between 20% and 350%. We further calculate the two-photon absorption cross-section and quantum yield of dopamine nanosensors, and confirm a two-photon power law relationship for the nanosensor excitation process. Finally, we show improved image quality of the nanosensors embedded 2 mm deep into a brain-mimetic tissue phantom, whereby one-photon excitation yields 42% scattering, in contrast to 4% scattering when the same object is imaged under two-photon excitation. Our approach overcomes traditional limitations in deep-tissue fluorescence microscopy, and can enable neurotransmitter imaging in the biologically-relevant milieu of the intact and living brain.

## 1. Introduction

Despite progress towards understanding brain function at a molecular and cellular level, insight into the detailed operation of brain circuits at an emergent level remains elusive.^[1]^ In particular, the development of optical sensors capable of monitoring *in vivo* neurotransmission with high spatial and temporal resolution can enable neuroscientists to broadly monitor the behavior of neuronal cells throughout neural circuits. Recently, semiconducting single-walled carbon nanotubes (SWNTs), which consist of a monolayer of graphene rolled into a cylinder with nanometer diameters and high aspect ratio, have emerged as an engineered nanomaterial readily adaptable for use in neuroscience and general bioimaging.^[2]^ The suitability of SWNTs to these applications arises from their small size (~10^−9^ m diameter, ~10^−6^ m length) and their inherent fluorescence that falls within the near infrared window (NIR-I, 700-950 nm; NIR-II, 950-1700 nm), coinciding with local minima in the absorbance and scattering spectra of water, blood and brain tissue, respectively.^[3]^ *In vivo* deep tissue imaging could be accomplished in the near infrared optical regime, in particular for transcranial detection of modulatory neurotransmitters whose biologically-relevant milieu is in the intact brain of awake and behaving animals. Recently, SWNT suspensions have been demonstrated as contrast agents for transcranial imaging of the mouse vasculature without the need for a cranial imaging window.^[4]^ In parallel, the utility of SWNT fluorophores was expanded beyond their use as a fluorescence contrast agent by functionalizing them with bio-mimetic polymers to recognize biological analytes – thus enabling their use as biosensors.^[5]^ To date, these fluorescent nanosensors have been used to detect a variety of biomolecules including dopamine, fibrinogen, nitric oxide, riboflavin and estradiol, to name a few.^[5–6]^ These *in vitro* nanosensors will likely engender real-time *in vivo* neuroscience imaging applications exploiting the biocompatibility, NIR emission, and remarkable photostability of functionalized SWNTs.

One major factor limiting fluorescence brain imaging has been signal attenuation from absorption and scattering of excitation photons. To mitigate these obstacles, neuroscientists and biologists have long relied on two-photon microscopy (2PM) using deep-penetrating, NIR light for the non-linear excitation of fluorophores.^[7]^ In neuroscience, 2PM has been employed to image strongly scattering brain tissue to study synaptic plasticity in cortical circuits,^[8]^ and to image the dynamics of Na^+^ and Ca^2+^ in intact central neurons in their native environment.^[9]^ The benefits of non-linear excitation imaging are also exploited in orthogonal fields of study, including tumorigenesis,^[10]^ embryogenesis^[11]^ and immunology.^[7c, 12]^ However, the photons emitted by two-photon excitation (2PE) are subject to the same absorption and scattering as those that emanate from one-photon excitation (1PE). Absorption and scattering of the signal from visible-wavelength photons emitted by fluorophores limit imaging depth^[13]^ and motivate the use of far-red and NIR fluorophores in brain imaging.^[14]^

The need for robust NIR fluorophores for multiphoton imaging, and a parallel interest in SWNTs for imaging and sensing applications, warrant an examination of nonlinear SWNT fluorescence for imaging. Multiphoton fluorescence imaging using NIR light has been demonstrated for a variety of different nanomaterials for biological imaging, but all suffer from the same drawback as many classical fluorophores: emission is confined to the visible or NIR-I windows.^[15]^ In contrast, non-linear excitation of SWNT nanosensors has the advantages of excitation *and* fluorescence emission in the NIR-II window,^[16]^ making them particularly well suited for imaging and sensing in brain tissue. Although SWNTs can be polymer-functionalized for use as real-time neurotransmitter nanosensors for potential *in vivo* imaging,^[6a]^ little work has been done to explore NIR 2PE of these functionalized SWNT nanosensors.

Here, we show that 2PE of both DNA-wrapped multi-chirality SWNTs, and surfactant-dispersed single-chirality SWNTs, is achievable with a 1560 nm femtosecond pulsed erbium laser. We compute the quantum yield and the two-photon absorption cross-section using (6,5) chirality SWNTs suspended in a sodium dodecyl sulfate (SDS) solution using a reference dye, 3,3′-diethylthiatri carbocyanine perchlorate (DTTC), whose quantum yield and two-photon cross section are known.^[17]^ Furthermore, we demonstrate molecular recognition of dopamine using our nanosensors and 2PE, achieving a two-fold increase in SWNT nanosensor fluorescence in the presence of 100 µM dopamine. Our results confirm that the molecular recognition principle is unaltered by the method of photoexcitation (1PE vs. 2PE), and provide a quantitative estimate of SWNT 2PE absorption cross section and quantum yield. Lastly, we show that 2PE yields significantly improved fluorescence spatial resolution over 1PE when SWNT are imaged 2 mm-deep in a strongly scattering Intralipid tissue phantom, motivating future *in vivo* applications in 2PM of nanosensors with near infrared fluorescence, henceforth referred to as dual NIR excitation-emission (NIR-EE) microscopy.

## 2. Results and Discussion

Motivated by the potential to use NIR SWNT neurotransmitter nanosensors for deep-tissue imaging, we first consider the relationship between imaging wavelength, imaging depth, and fluorescence attenuation, to identify minimum attenuation wavelengths for excitation and emission. Drawing from literature values^[4a]^ for wavelength dependent absorbance of adult mouse scalp skin (1 mm), cranial bone (1 mm), and water, and scattering coefficient values^[4a]^ for scalp skin, cranial bone and brain tissue, we calculate the total wavelength dependent optical density as the sum of absorption and scattering as a function of depth (Figure 1a). Figure 1a shows that SWNT NIR emission coincides with a local attenuation minimum in the 1000-1400 nm range, adjacent to a local water absorption peak at 1400 nm. Figure 1a also shows a second attenuation minimum in the 1600-1800 nm, which we identify as the optimal 2PE window. Thus, to evaluate SWNTs as NIR fluorophores for two-photon microscopy, we compared fluorescence emission spectra of aqueous solutions of well-dispersed SWNTs using 1PE and 2PE. Using a HeNe laser (633 nm) as a 1PE source, and a femtosecond erbium laser (1560 nm, near the second attenuation minimum) as a 2PE source, we collected emission spectra for two dispersed SWNT samples, i) A dopamine responsive nanosensor composed of HiPco SWNTs wrapped with the single stranded DNA sequence (GT)15 and ii) SWNTs suspended in SDS and enriched to contain primarily (6,5) chirality SWNTs. The 2PE source wavelength (1560 nm) is far from the linear absorbance of SWNTs (500-900 nm), ensuring that any fluorescence emission observed is on account of non-linear absorption processes. For both 1PE and 2PE, we employed a perpendicular-geometry fluorescence excitation/detection setup (Figure 1b) with a quartz cuvette mounted on a pair of translation stages that enabled accurate placement of the fluorophore sample with respect to the pump-beam waist, for spectroscopy and imaging experiments. We note that an upright or inverted microscopy setup will be most suitable for imaging experiments alone. The pump light was focused into the cuvette containing SWNT or a reference dye, (DTTC), with a waist radius of 5 µm and a confocal distance of 100 µm in free space. Fluorescence emission from the focal volume was collected at 90 degrees and imaged onto a Princeton Instruments SCT 320 spectrometer with a liquid nitrogen cooled InGaAs array (Figure 1b). Details of the optical setup are outlined in the Supporting Information.

**Figure 1.**
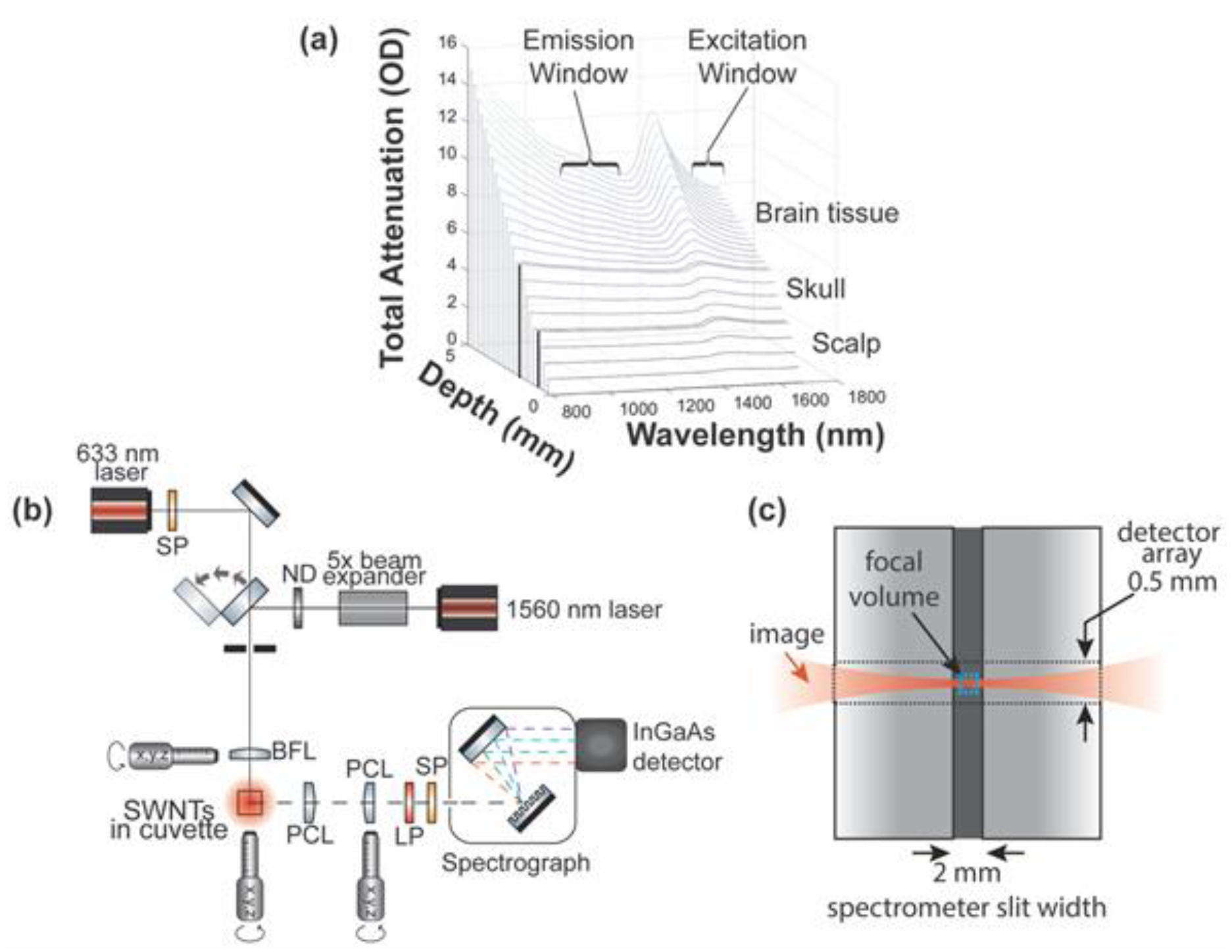
NIR-EE fluorescence microscopy for imaging in highly scattering media. (a) Total photon attenuation (absorbance and scattering) for 1 mm layers of scalp and skull as a function of imaging depth into mouse brain tissue. Both emission and 2-photon excitation of SWNTs fall within local attenuation minima. (b) Schematic depicting collection of fluorescence emission spectra from SWNT samples using the 633 nm laser for 1PE and 1560 nm laser for 2PE. BFL: best-form lens, PCL: planoconvex lens, SP: 1500 nm shortpass filter, LP: 860 nm longpass filter. Light path terminates into a spectrometer with an InGaAs detector. (c) Magnified diagram showing the focal volume of the laser in the cuvette relative to the spectrometer slit and detector array height.

As expected, the fluorescence emission spectra of multi-chirality (GT)15DNA-wrapped SWNT dopamine nanosensors using 1PE contains multiple peaks in the NIR-II range corresponding to the convolution of emissions from a mixture of different SWNT chiralities (Figure 2a).^[18]^ We deconvolve the contributions of each SWNT chirality to emission peaks using a non-linear least squares method (See Supporting Information **Figure S1** and **S2**, **Table S1** and **S2**).^[5]^ The prominent emission peaks at 1030 nm and 1126 nm are consistent with on-resonance excitation of (7,5) and (7,6) chirality SWNTs, respectively, in addition to contributions from other chiralities excited off-resonance. The emission peak at 1265 nm is a convolution of (10,3), (9,5) and (11,1) chirality SWNTs excited on-resonance, along with off-resonance SWNTs. Emission peaks from additional SWNT chiralities excited off-resonance are also visible.

**Figure 2.**
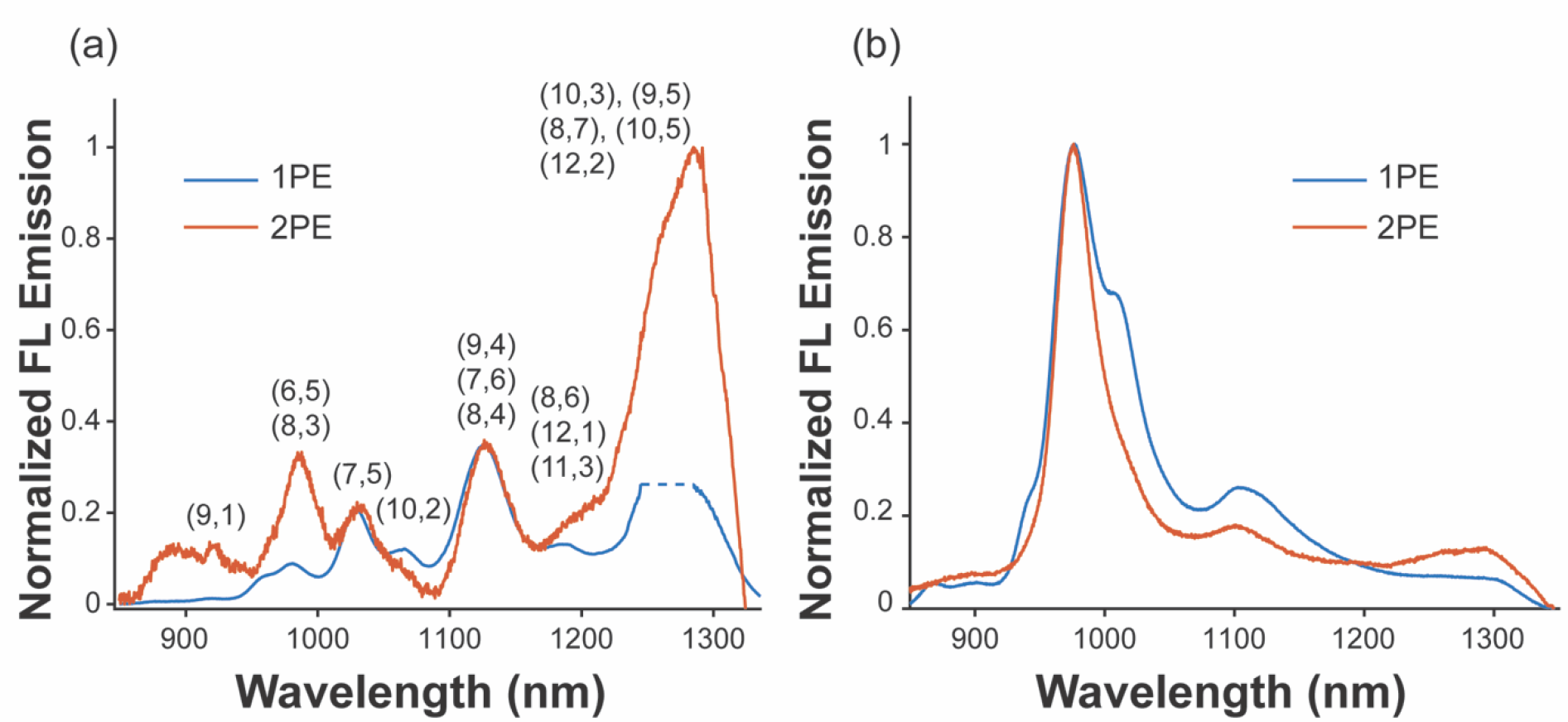
1PE and 2PE emission spectra of SWNT. Fluorescence emission of (a) (GT)15DNA-SWNT and (b) SDS-(6,5)-chirality SWNT suspensions excited with a 633 nm CW 1 photon excitation source (blue) or a 1550 nm fs-pulsed 2 photon excitation source (orange). Labels indicate the chiralities that contribute to each peak. Note: dotted line indicates region of detector saturation caused by 2^nd^ order scattered laser light.

Using 2PE, we observed comparable fluorescence emission from (GT)15DNA-SWNT nanosensors to establish that SWNT-based dopamine nanosensors are amenable to multi-photon excitation in aqueous solution (Figure 2a). As expected, the 2PE emission spectrum of the DNA-wrapped SWNTs reveals a change in the chiralities excited on-resonance when compared to the 1PE emission spectrum. Normalizing both 1PE and 2PE (6,5) chirality SWNT emission spectra to the 1126 nm peak (dominant peak for 1PE) shows a three-fold increase in relative emission intensity, indicating an increase in SWNT excitation efficiency. Our measurements are consistent with previous reports of 2PE emission for (6,5) chirality SWNT, which show on-resonance excitation near 1560 nm for dried or surfactant suspended samples.^[16, 19]^ A similar increase in excitation efficiency is observed for chiralities emitting above 1200 nm, which has not been previously reported. This increase likely arises from near-resonant 2PE excitation of (10,3) and (11,1) chiralities, which share 1PE emission peaks closer to that of (6,5) SWNTs.

To further characterize 2PE emission of SWNTs, we prepared a sample enriched with (6,5) chirality dispersed SWNTs through previously reported protocols.^[20]^ Briefly, HiPco SWNT dispersed in SDS by probe tip sonication were purified by adsorption column chromatography through separation over a series of sephacryl gel beds. Through this process, we obtain a chirality-purified fraction of enriched (6,5) SWNT, while eliminating SWNT bundles, aggregates and non-emissive metallic SWNTs. Absorbance spectrum measurements of the purified sample (Supporting Information **Figure S3a**) reveal prominent peaks associated with the *E22 and E11* transitions of (6,5) chirality SWNTs. The 1PE emission spectrum (Figure 2b) shows emission peaked at 975 nm, corresponding to (6,5) chirality SWNTs, with additional fluorescence peaks near 1000 nm and 1100 nm resulting from other SWNT chiralities remaining after purification. The relative emission intensity of these off-resonance chiralities, however, is minimal. 2PE excitation of the (6,5) chirality was tested by changing the laser excitation source from a 633 nm CW to the 1560 nm fs-pulsed erbium laser. 2PE emission spectra from (6,5) SWNT show a dominant emission peak at 975 nm (Figure 2b), as with 1PE excitation. Furthermore, the fluorescent contribution of non-(6,5) chiralities decreases in the spectrum collected using 2PE compared to 1PE due to off-resonance excitation, consistent with our observations of the mixed-chirality SWNT sample.

Emission intensity of the (6,5) chirality SWNTs and (GT)15-DNA SWNTs increased without a change in the emission lineshape as the 2PE laser power was increased (Figure 3a, 3c). The integrated emission intensity as a function of laser power exhibited a quadratic dependence as indicated by a slope of approximately 2 for the SWNT fluorescence intensity versus laser excitation power plotted on a log-log scale (Figure 3c, 3d), confirming the SWNT fluorescence resulted from a non-linear photon absorption process.

**Figure 3.**
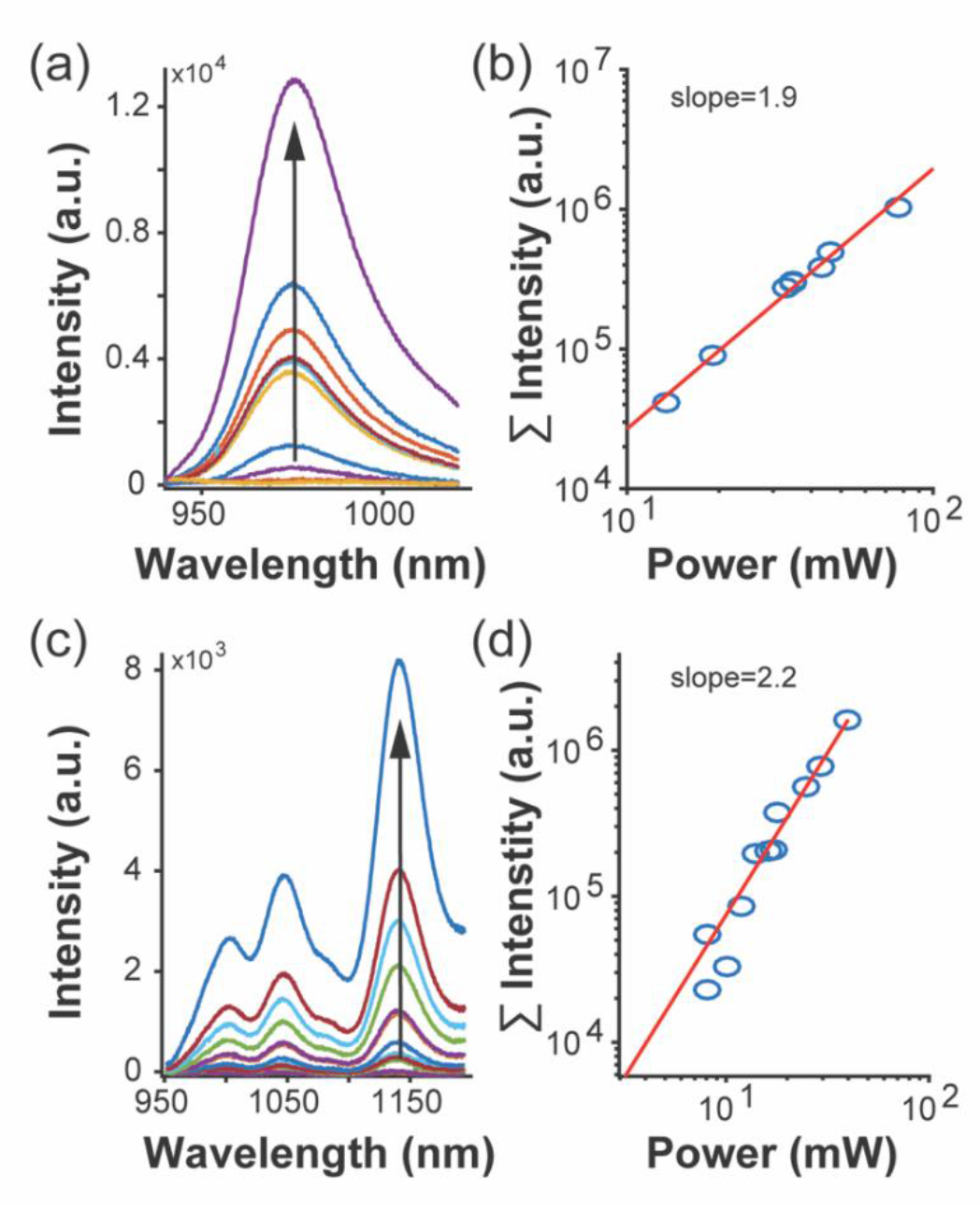
Emission spectra of SDS-(6,5)-enriched SWNTs and GT15-DNA SWNTs excited by two-photon excitation at different laser powers. The integrated emission intensity increases quadratically with laser power as shown by the near 2 slope of a log-log plot of the emission intensity as a function of laser power for both (a)-(b) SDS-(6,5)-enriched SWNTs and (c)-(d) GT15-DNA SWNTs. A background signal, comprised of the integrated intensity at an incident power below which appreciable changes in signal were observed, was subtracted prior to integration.

The 1PE quantum yield, ***Q*_1*P*_**, and absorption cross section for 2PE, ***σ*_2*P*_**, are useful values for comparing fluorophores for use in 2PM, as the fluorescence signal from 2PE depends linearly on these two values (see Supporting Information for details). ***Q*_1*P*_**, can be calculated for a sample (*S*) fluorophore by comparing its behavior to a reference (*R*) fluorophore using:^[21]^

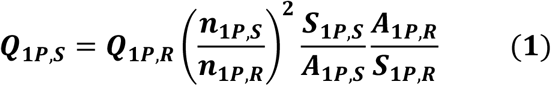

where ***S*** is the number of emission photons detected per unit time, ***A*** is the absorbed fraction of the impinging pump light, ***n*** is index of refraction, and subscripts *S* and *R* represent the sample fluorophore and reference fluorophore, respectively. ***S*** is measured by the integral:

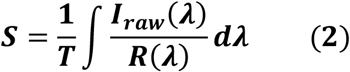

where ***T*** is the integration time period, ***I_raw_(λ)*** is the baseline-corrected raw intensity of detected emission photons, and ***R(λ)*** is the wavelength-dependent response of the spectrometer and dark current offsets. With ***Q*_1*P,S*_** calculated, ***σ*_2*P,S*_** can be computed using a similar comparison:^[21]^

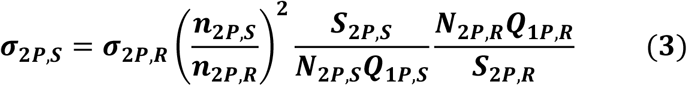

where ***N*** is the number density of fluorophores per unit volume. Note that errors in ***Q*_1*P,S*_** propagate into the ***σ*_2*P,S*_** estimate. It is assumed that pumping is below the onset of saturation for both the 1PE and 2PE systems.

For this work, SWNT cross section measurements were made for purified (6,5) SWNT because the well-characterized extinction coefficient of chirality-purified SWNT is necessary for the calculation of ***Q*_1*P,S*_** (see Supporting Information for details). This was in lieu of the multi-chirality, and thus multi-emitter, suspension of GT15-SWNT nanosensors, each with a unique extinction coefficient, which would confound ***Q*_1*P,S*_** measurements. SWNT absorption cross-section is invariant upon exposure to dopamine, thus the cross section of purified (6,5) SWNTs provides a reasonable order of magnitude estimate of ***σ*_2*P,S*_** for SWNTs of various wrappings and chiralities. The reference fluorophore used was 3,3′-diethylthiatricarbocyanine perchlorate (DTTC), as it is a NIR emitting dye with previously characterized ***Q*_1*P*_** and ***σ*_2*P*_**.^[17]^ Absorbance and emission spectra are shown in the Supporting Information (**Figure S3, S4** and **S5**). Calibrations and additional considerations for ***Q*_1*P*_** and ***σ*_2*P*_** calculations as well as calibration-corrected spectra are outlined in the Supporting Information (**Figure S6, S7, S8** and **S9**). Following these methods and calculations we derive ***Q*_1*P,SWNT*_** = **0.0023** and ***σ*_2*P,SWNT*_** = **239,000** GM. Note that 1 GM, the unit of two-photon absorption cross section, is defined as 10^−50^ cm^4^ sec photon^**−**1^. Values for ***Q*_1*P,SWNT*_** reported in literature range from 10^**−**4^ to a few percent, whereas ***σ*_2*P,SWNT*_** can be inferred from previous work as ranging from 10,000 GM to 700,000 GM.^[16, 22]^ Both our ***Q*_1*P,SWNT*_** and ***σ*_2*P,SWNT*_** fall within ranges expected based on data from these previous reports. A detailed list of input values is included in **Table S3** and **S4** of the Supporting Information. Given our experimental conditions, we would expect an imaging resolution of approximately the beam waist, 4.8 µm, to produce an image using pixel dwell times on the order of milliseconds. Micron length and millisecond time frames are in-line with the requisite imaging parameters necessary to capture relevant processes in brain neurotransmission.

Next, we evaluated the efficacy of 2PE for neurotransmitter detection using a dopamine-sensitive SWNT nanosensor. (GT)_15_DNA polymer-functionalized SWNTs are recently discovered NIR optical nanosensor enabling the selective and reversible sensing of the neuromodulatory neurotransmitter dopamine.^[6a]^ These SWNT-based dopamine nanosensors exhibit a marked increase in fluorescence emission intensity upon binding dopamine with sensitivities down to 10 nM. To date, the fluorescence response of SWNT dopamine nanosensors has only been characterized using one-photon visible wavelength excitation. We aim to determine whether SWNT-based dopamine nanosensors are compatible with NIR-EE microscopy for multi-photon deep-tissue imaging applications, which requires the nanosensor’s ‘turn-on’ fluorescence response to dopamine to be indifferent to the SWNT photon absorption process. Using both 1PE and 2PE, fluorescence emission spectra were collected from a sample of (GT)_15_DNA-SWNT dopamine nanosensors before and after the addition of 100 µM dopamine (Figure 4a). Adding dopamine increases the fluorescence emission intensity for all SWNT chiralities for both 1PE and 2PE (Figure 4b and 4d), confirming that the response is independent of the photon absorption process. We calculated the relative change in peak fluorescence intensity for different SWNT chiralities by first deconvolving the SWNT emission spectra (see Supporting Information) and calculating the dopamine nanosensor signal, (*I-I*_0_)/*I*_0_, where *I*_0_ is the integrated fluorescence intensity for each SWNT chirality before dopamine addition. We combined the integrated intensities into groups of chiralities that had indistinguishable emission spectra, e.g. the integrated emission of (9,1), (8,3), and (6,5) were combined into a single group. For both excitation methods, the fluorescence intensity of SWNT nanosensors increases with the addition of dopamine for all SWNT chiralities. For 1PE (Figure 4b and 4c), the response is only mildly dependent on chirality and varies between 145% and 255%, comparable to previous measurements.^[6a, 23]^ The fluorescence response using 2PE (Figure 4d and 4e) shows considerably more chirality dependence, with maximal dopamine-induced fluorescence increases observed for longer NIR-II wavelengths (1100 nm – 1350 nm). The chirality dependence observed for 2PE is likely due to differences between the excitation efficiencies of different chirality SWNTs using NIR compared to visible light. The benefit of 2PE excitation of dopamine nanosensors, combined with their enhanced NIR-II emissions above 1100 nm (Figure 1a), motivates their use in highly scattering biological media and for deep-tissue imaging using 2PM for NIR-EE microscopy.

**Figure 4.**
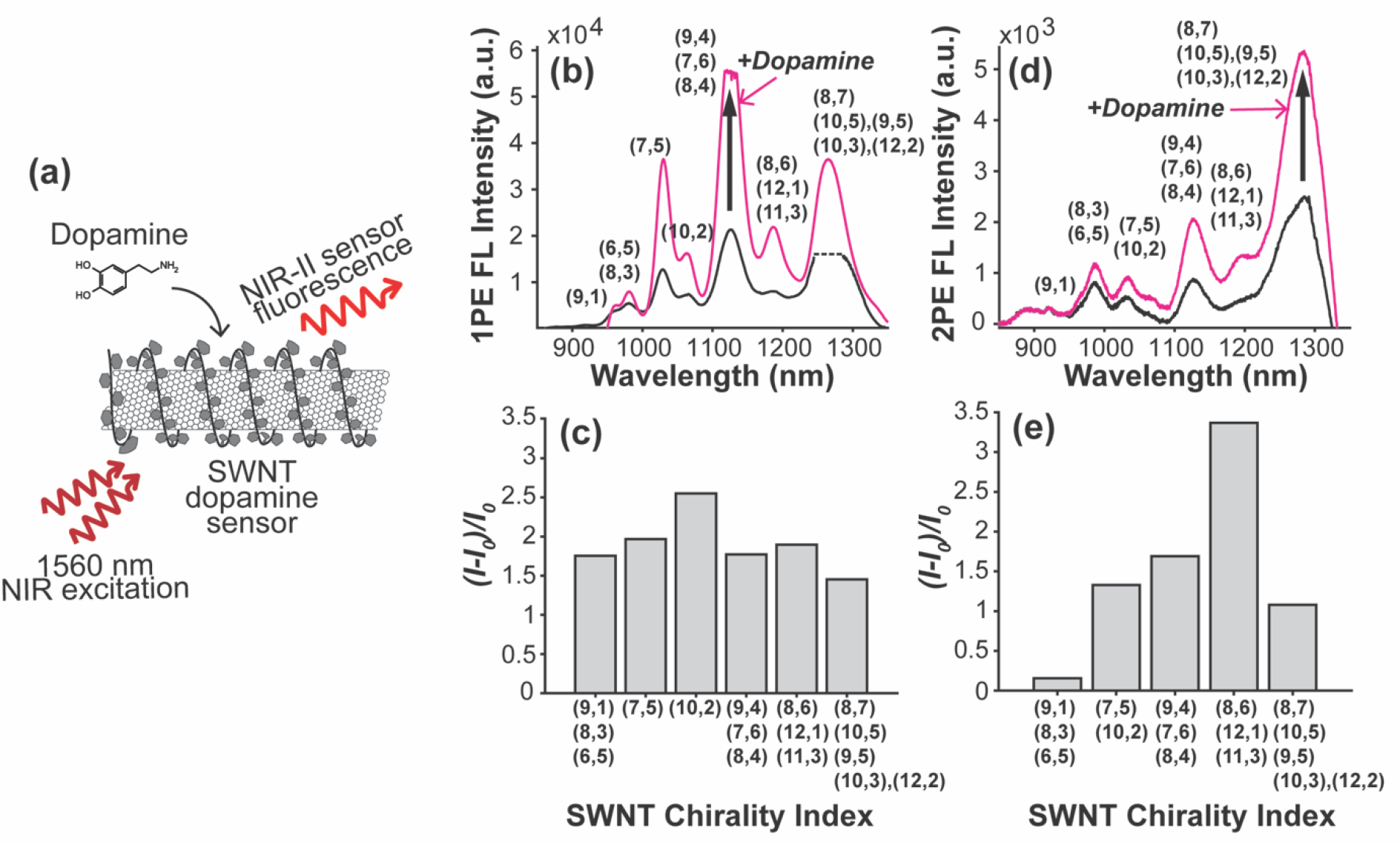
Fluorescence response of dopamine nanosensors using one-and two-photon excitation. (a) Schematic depicting dopamine binding a dopamine nanosensor and the resulting fluorescence emission enhancement. Dopamine (100µM) increases the fluorescence emission intensity of (GT)_15_DNA wrapped SWNTs using both 1PE (b)-(c) and 2PE (d)-(e). Note: dotted line indicates region of detector saturation caused by 2^nd^ order scattered laser light.

To demonstrate the advantages of two-photon over one-photon fluorescence imaging of SWNT-based nanosensors, we compared reconstructed images of a SWNT-filled capillary immersed in a highly scattering tissue phantom, 1% Intralipid solution,^[3c, 4a, 13]^ under 1PE versus 2PE. Of relevance to deep-brain imaging of dopamine, the tissue phantom simulates both the absorbance due to water and the scattering in brain tissue.^[4a]^ An 800 µm inner-diameter quartz capillary was filled with a (6,5)-chirality purified SWNT or (GT)_15_-DNA SWNT suspension and placed into the cuvette containing 1% Intralipid (see **Experiment Section**) at an Intralipid depth of 0.5 mm from the surface of the cuvette facing the excitation focusing lens (excitation Intralipid depth), and an Intralipid depth of 2 mm from the capillary surface facing the InGaAs detector (emission Intralipid depth). We reconstruct a 1-dimensional image of the capillary by scanning the capillary laterally across the focused beam using a translation stage and collecting the emitted NIR fluorescence in the transverse direction (Figure 5a). Each pixel of this one-dimensional line scan is generated by the summed detector intensity at each focusing lens position (Figure 5b–5c), and used to reconstruct an image of the capillary (Figure 5d and 5e). The scattering of 633 nm excitation photons by the Intralipid occluding the intra-capillary SWNTs produces a severely blurred image (Figure 5d–5e, **left**) with poorly resolved edges relative to the true capillary edges. Scattering of excitation photons against scattering media causes photons to deviate from the trajectory of their laser source, such that they excite fluorophores outside of the focal volume. As such, the reconstructed image includes SWNTs excited outside of the focal volume when using 1PE. Visible wavelength excitation scattering occurred even when the laser focus was as far as 1 mm from the true edge of the capillary. In contrast, the image constructed using 2PE via NIR pulsed laser excitation resolved sharp edges of the capillary (Figure 5d–5e, **right**), allowing for precise image reconstruction of the capillary diameter. Bright peaks outside the capillary region are a result of fluorescence generated by pump light scattered off the outer capillary edges.

**Figure 5.**
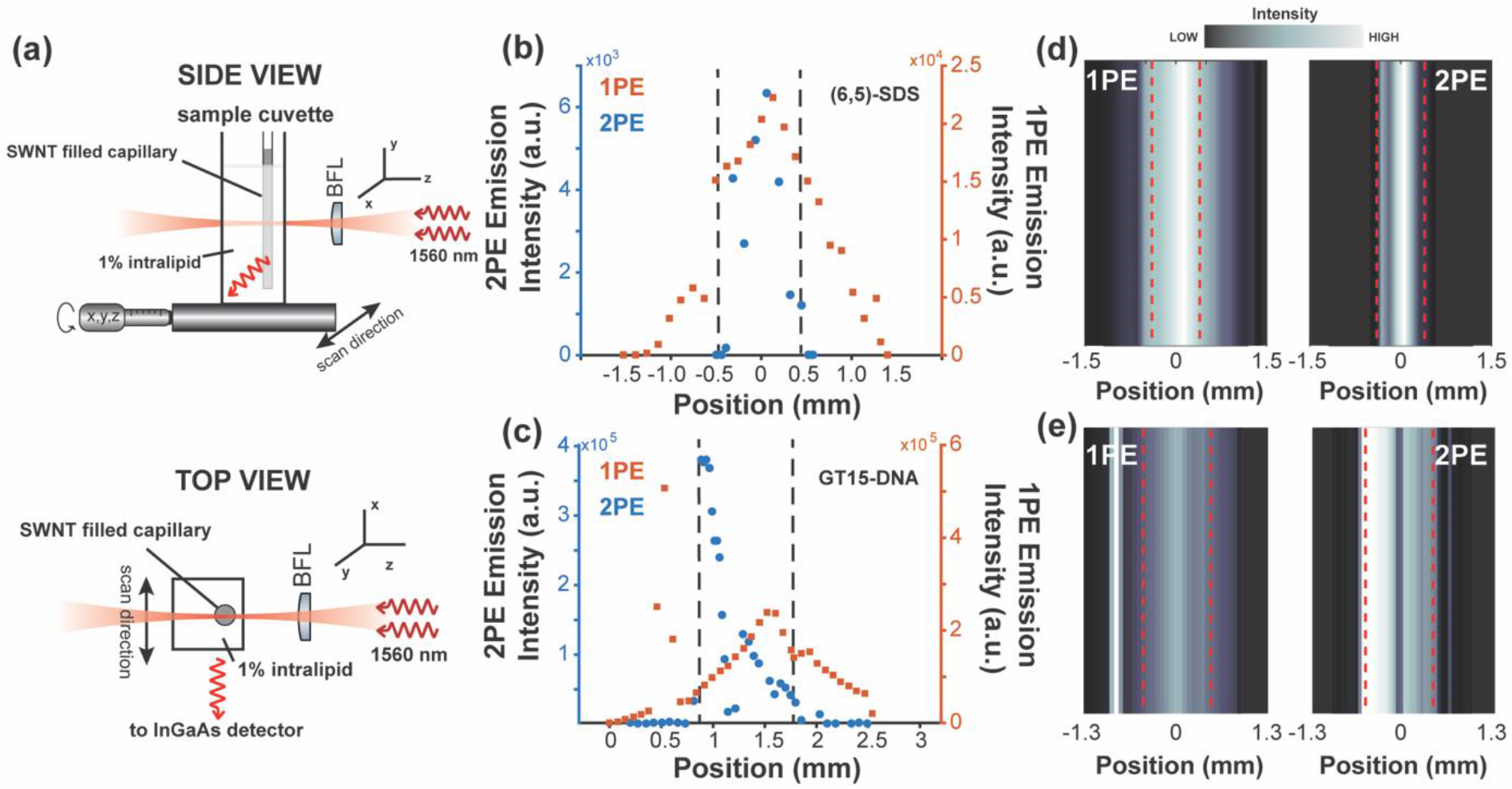
Improved spatial resolution at 2 mm Intralipid imaging depth with NIR-EE microscopy. (a) Schematic depicting a SWNT-filled 0.8 mm-inner diameter capillary submerged in a strongly scattering solution (1% Intralipid) at a depth of 0.5 mm from the excitation edge of the cuvette and at an Intralipid depth of 2 mm from the cuvette emission edge. The sample cuvette and capillary were scanned in the x-direction through the beam focus. Integrated fluorescence intensity as a function of scan position as the capillary (filled with (a) (6,5)-purified SWNTs and (b) (GT)_15_-DNA wrapped SWNTs) is scanned across the focused beam using 1PE (red squares) and 2PE (blue circles). For reference, the dashed black lines indicate the position of the capillary inner edge. The 1000 nm – 1300 nm fluorescence intensity data presented in (b) and (c) is reconstructed into one-dimensional line scan images for (d) (6,5)-purified SWNTs and (e) (GT)_15_-DNA wrapped SWNTS. Dashed red lines indicate the position of the capillary inner edge.

Further evidence for the advantages of NIR 2PE for imaging is found by examining images of the pump beam in a highly scattering environment comprised of (GT)_15_-DNA wrapped SWNTs diluted in a 1% Intralipid solution. An image of the 1PE pump beam generated by a combination of SWNT fluorescence and scattered pump light (Figure 6a) shows a greater degree of scattering than the 2PE pump beam (Figure 6b). Additionally, the 2PE beam penetrates to depths of approximately 2 mm into the highly scattering solution.

**Figure 6.**
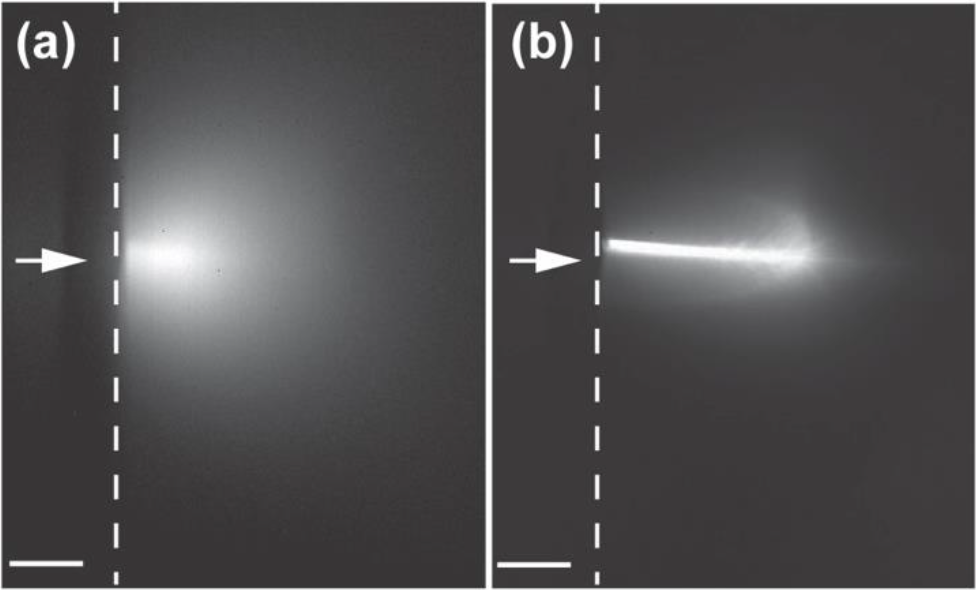
Scattering of pump beam in scattering media. Images generated using light collected between 900 - 1300 nm at the focus of the 1PE pump beam (a) and 2PE pump beam (b) in a solution containing 1% Intralipid and 10 mg/L (GT)_15_-DNA SWNTs. Arrows indicate direction of beam propagation and dotted lines indicate the quartz/sample interface at the inner edge of the cuvette. Scale bar is 1 mm.

These results demonstrate the advantages of using 2PE for deep-tissue imaging in combination with NIR-I and NIR-II emissive fluorophores, namely reduced scattering of incident excitation photons. Herein, we show that NIR-EE microscopy builds upon the advantages of conventional 2PE by both exciting and collecting photons in local tissue transparency minima (Figure 1a). By minimizing scattering of incident photons, we can enhance localization of excitation to improve imaging quality. For our capillary images, up to 42% of the integrated collected fluorescence occurred beyond the extent of the physical capillary boundaries using 1PE compared to only 4% scattering-induced image blurring using 2PE. Additionally, bio-mimetic polymer functionalization of SWNT for molecular detection of modulatory neurotransmitters expands their utility beyond biological contrast imaging agents, and demonstrates their inherent advantages for detection of biological targets in strongly scattering tissues. In particular, NIR-EE imaging of SWNT-based modulatory neurotransmitter nanosensors could enable real-time fluorescence monitoring of analytes such as dopamine in optically dense brain tissue.

## 3. Conclusion

In summary, we demonstrate NIR-II fluorescence-based imaging of the neurotransmitter dopamine using NIR-II laser excitation of functionalized SWNT nanosensors. To evaluate the efficacy of using 2PE of SWNT nanosensors for imaging applications, we measure the quantum efficiency, ***Q*_1*P*_**, and absorption cross section of 2PE, ***σ*_2*P*_**, as 0.0023 and 239,000 GM, respectively. Using the current configuration, these values establish reasonable estimates for laser dwell times on the order of milliseconds for constructing images with resolution of ~5 µm using scanning imaging microscopy. Comparisons of 1PE and 2PE scanning imaging of a NIR SWNT-filled capillary in turbid Intralipid tissue phantoms confirm that the reduced scattering of NIR-II incident excitation light improves fluorescence spatial resolution and imaging quality as shown by sharper image boundaries and better localization of integrated SWNT NIR fluorescence. Our results inspire the use of NIR SWNT nanosensors and 2PE in future investigations involving real-time imaging of dopamine directly in the highly-scattering tissue of the brain.

## 4. Experimental Section

### Imaging setup and alignment

One-photon pump light was generated using a ~5 mW CW 633 nm He-Ne laser with a 1000 nm shortpass cleanup filter and with a 1 mm beam radius at the focusing lens. 2PE pump light was generated using a single-mode-fiber-pigtailed pulsed Erbium laser (ELMO-HIGH POWER, Menlo Systems) with nominal wavelength, repetition rate, and pulsewidth of 1560 nm ± 30 nm, 100 MHz, and <90 fsec respectively. The measured beam power at the cuvette after collimation and 5X beam expansion was 77 mW with a beam waist of 4.1 mm. For measurements of fluorescence as a function of laser power for (GT)_15_-SWNTS (Figure 3c–3d), capillary imaging experiments using (GT)_15_-SWNTS (Figure 5c–5d) and beam imaging (Figure 6), a comparable pumped laser source was used (IMRA femtolite F-100) with a nominal wavelength, maximum beam power at cuvette, and waist of 1590 nm ± 10 nm, 145 mW and 4.1 mm, respectively. The pump light was focused with a 40 mm focal-length “best form” lens into the cuvette containing DTTC or SWNT, from which fluorescence emission was imaged onto a Princeton Instruments SCT 320 spectrometer slit with a pair of 50 mm focal-length plano-convex lenses of 25 mm diameter. Images of the scattered pump beam (Figure 6) were obtained using a Princeton Instruments NIRvana InGaAs camera in place of the spectrometer. Protected silver coated mirrors (Thorlabs) were used to direct the beam. An 860 nm longpass filter, 1500 nm shortpass filter and an RG9 colored glass filter (Thorlabs) were used to exclude excitation light from the spectrometer. A 1400 nm longpass filter (Thorlabs) was used to clean up the pump beam. Accurate height alignment was needed to center the image of the fluorescence stripe on the slit region whose wavelength-dispersed image was in turn relayed to the PyLon linear detector with a 0.5 mm pixel height. Using a 150 groove mm^−1^ diffraction grating, the 1024-element detector array (with 25 µm pixel pitch and ~25 mm length) covered a wavelength region of ~500 nm. The resultant ~20 nm mm^−1^ dispersion, coupled with typical ~1 – 2 mm slit widths, yielded spectral resolutions on the order of tens of nm.

### DTTC and SWNT sample preparations

Suspensions of (GT)_15_-DNA SWNT dopamine nanosensors were prepared by adding ~1 mg of dried raw HiPco SWNTs (NanoIntegris) to a 1 mL solution of 100 µM ssDNA (Integrated DNA Technologies, standard desalting) in 100 mM sodium phosphate buffer (pH 7.4). The solution was then sonicated for 5-10 minutes in a bath sonicator followed by 10 minutes of probe tip sonication (3 mm tip diameter (CV18, Ultrasonic Processor, Cole Palmer). The sample was then centrifuged for 30 minutes at 16,100 × g to pellet unsuspended SWNTs, aggregates and bundles, while keeping the supernatant. The concentration of suspended SWNTs was estimated by absorbance measurements at 632 nm using an extinction coefficient of 0.036 L cm^−1^ mg^−1^. Fluorescence emission spectra (1PE and 2PE) were collected from a solution diluted with 1X PBS buffer (137 mM NaCl, 2.7 mM KCl, 10 mM Na2HPO4, 1.8 mM KH2PO4, using MilliQ Millipore deionized water) to a final concentration of 10 mg L^−1^.

Solutions of (6,5)-enriched SWNTs were prepared as described previously.^[20]^ Raw HiPco SWNT were sonicated in 2% SDS for 20 hours followed by 4 hours of centrifugation at 187,000 × g, keeping the top 90% of the supernatant. Sodium cholate (SC) was then added to the solution to a final concentration between 0.1 and 0.7%. SWNTs solutions were then passed through columns containing Sephacryl 200 gel. Columns were then rinsed with 175 mM SDS and collected into fractions, which were then characterized by absorption spectroscopy. The final concentration of (6,5)-enriched SWNTs was measured to be 3.6 × 10^−7^ M (see Supporting Information for details).

The spectral contribution from nanotubes of different chirality was deconvolved using a custom written script using MATLAB as described previously.^[5]^ Chiralities with significantly overlapping emission spectra were grouped together by integrating their intensity contributions as calculated by non-linear least squares optimization of FWHM, peak area, and center wavelength to fit a Lorentz distribution. Output of optimized parameters and resulting chirality distributions for our samples are included in the Supplementary Information **Table S1-S2, S5-S6**. The slopes of fluorescence intensity plotted against laser power were calculated using a nonlinear least squares fit of a power function (power1) using MATLAB’s fit() function (MATLAB 2016a, The MathWorks).

### Dopamine nanosensor NIRF response assay

To measure the fluorescent response of SWNT nanosensors to dopamine, the NIR emission spectra were collected from a sample cuvette containing 10 mg L^−1^ (GT)_15_DNA SWNT dopamine nanosensors in 1X PBS buffer. A concentrated solution of freshly prepared dopamine-HCl (Sigma) in deionized water was added to the cuvette containing 10 mg L^−1^ (GT)_15_DNA SWNTs to a final concentration of 100 µM, and allowed to incubate for 5 minutes prior to collecting NIRF emission spectra.

### Intralipid sample preparation

A 0.8 mm inner diameter capillary tube (PYREX, 0.8-1.1×100 mm) was filled with a SWNT nanotube suspension using capillary forces and sealed using 1% agarose gel. The capillary was suspended in the cuvette containing a solution of 1% Intralipid (Intralipid, 20% emulsion, Sigma Life Sciences) in deionized water using a V-mount attached to a linear translation stage for precise positioning. The cuvette was positioned 0.5 mm from the cuvette edge facing the pump source and 2 mm from the cuvette edge facing the spectrometer as shown in Figure 5b in the main text. The position of the objective lens was adjusted to place the beam focus at the same depth as the capillary. The entire cuvette/capillary assembly was scanned laterally across the beam focus using a translation stage in increments of 0.005 inches. The total fluorescence intensity from the sample in the capillary was collected at each position by integrating across the entire sensor array of the spectrometer.

## Supporting Information

Supporting Information is available from the Wiley Online Library or from the author.

## Acknowledgements

Our work was supported by a Burroughs Wellcome Fund Career Award at the Scientific Interface (CASI), the Simons Foundation, a Stanley Fahn PDF Junior Faculty Grant with Award # PF-JFA-1760, and a Beckman Foundation Young Investigator Award (M.P.L.). A.B. Acknowledges the support of an NSF Graduate Research Fellowship. We gratefully acknowledge technical guidance and equipment from Simon Kocur and Menlo Systems, Inc.

